# Comparison between traditional FFT and marginal spectra using the Hilbert-Huang transform method for the broadband spectral analysis of the EEG in healthy humans

**DOI:** 10.1101/2020.06.06.137950

**Authors:** Eduardo Arrufat-Pié, Mario Estévez Báez, José Mario Estévez Carreras, Calixto Machado Curbelo, Gerry Leisman, Carlos Beltrán

## Abstract

The fast Fourier transform (FFT), has been the main tool for the EEG spectral analysis (SPA). However, as the EEG dynamics shows nonlinear and non-stationary behavior, results using the FFT approach may result meaningless. A novel method has been developed for the analysis of nonlinear and non-stationary signals known as the Hilbert-Huang transform method. In this study we describe and compare the spectral analyses of the EEG using the traditional FFT approach with those calculated with the Hilbert marginal spectra (HMS) after decomposition of the EEG with a multivariate empirical mode decomposition algorithm. Segments of continuous 60-seconds EEG recorded from 19 leads of 47 healthy volunteers were studied. Although the spectral indices calculated for the explored EEG bands showed significant statistical differences for different leads and bands, a detailed analysis showed that for practical purposes both methods performed substantially similar. The HMS showed a reduction of the alpha activity (−5.64%), with increment in the beta-1 (+1.67%), and gamma (+1.38%) fast activity bands, and also an increment in the theta band (+2.14%), and in the delta (+0.45%) band, and vice versa for the FFT method. For the weighted mean frequencies insignificant mean differences (lower than 1Hz) were observed between both methods for the delta, theta, alpha, beta-1 and beta-2 bands, and only for the gamma band values for the HMS were 3 Hz higher than with the FFT method. The HMS may be considered a good alternative for the SPA of the EEG when nonlinearity or non-stationarity may be present.

## 1. Introduction

The electroencephalogram (EEG) represents the integral bioelectrical activity of the brain, recorded from the scalp, and is generated by the interaction of the whole neuronal activity. The EEG has been extensively used for the study of multiple conditions in animals and humans. The information carried by indices calculated in the frequency domain of the EEG has been the subject of attention for many years. Nonparametric methods that apply the fast Fourier transform (FFT), have been the main tool to obtain the spectra from which calculate the indices in the frequency domain of the EEG. However, the EEG dynamics shows nonlinearity and non-stationarity (Noshadi et al. 2014; Abdulhay et al. 2017; Alegre-Cortes et al. 2016; Soler et al. 2020) and strictly speaking, the FFT methods are limited to linear systems. Therefore, in some conditions the interpretation of the results obtained through the FFT can be meaningless. A novel fully data-driven method (Huang et al. 1998). for the analysis of non-Gaussian, nonlinear and non-stationary signals, the empirical mode decomposition (EMD), followed by the Hilbert transform of the extracted modal components with the EMD, known also as the Hilbert-Huang transform method, has been successfully used in geophysical studies (Huang et al. 1998; Huang, Wu 2008), image analysis (Nunes et al. 2003), thermal profiles analysis (Subhani et al. 2016), and power quality analysis (Camarena-Martinez et al. 2016). EMD has demonstrated itself to be a reliable and effective method in the processing of different biomedical signals such as the EEG (Al-Subari et al. 2015; Al-Subari et al. 2016; Amo et al. 2017; Estevez-Baez et al. 2017a, b; Hassan, Bhuiyan 2017; Hou et al. 2018; Rahman, Fattah 2017; Carella et al. 2018; Chatterjee 2019; Dinares-Ferran et al. 2018; Chuang et al. 2019; Hansen et al. 2019; Javed et al. 2019). There are two main procedures to calculate and graphically represent the results of the Hilbert-Huang method: the time frequency Hilbert spectrum, and as an alternative the Hilbert marginal spectrum (HMS), consisting in selecting a lower frequency resolution value, leaving the time axis undisturbed. The marginal spectrum offers a measure of total amplitude (or energy) contribution from each frequency value and its graphical representation is very similar to the spectrum calculated by other traditionally used methods including the FFT: The X-axis includes the spectral discrete frequencies, and the Y-axis the corresponding amplitude values (power).

The HMS has been used by different authors after applying the Hilbert-Huang method to the EEG signal (Chen et al. 2010; Li 2006; Li et al. 2008; Chen et al. 2016; Xiangjun et al. 2017; Zhu et al. 2015; Park et al. 2011) but to our knowledge, the possible differences between the results obtained using the HMS and the conventional FFT spectra in resting conditions in healthy humans have not been studied in detail, and it results imperative, because for many years the universally used method for the spectral analysis of the EEG has used the FFT spectra. The advantages and possible limitations of the novel alternative approach, in this case the use of the HMS, could be better appreciated and could contribute to their more extended use in EEG investigations. Therefore, in this study we aimed to describe and compare quantitative indices of the spectral analyses of the EEG calculated using the concepts of the empirical mode decomposition (EMD) and the marginal Hilbert spectra (MHS), with those calculated using the traditional approach of the fast Fourier transform (FFT), to analyze the origin of the possible differences in the estimation of the power spectral density (PSD), and finally to comment the results.

## 2. Methods

### 2.1 Participants and general experimental profile

Forty-seven healthy right-handed volunteers were included in this study. Demographic and some vital indices of the participants are shown in Table 1.

**Table 1.** Demographic data and some vital signs of the forty-seven healthy volunteers.

| Gender | n | Age<br>(m ± sd) [range] | RR<br>(m ± sd) | HR<br>(m ± sd) | SBP<br>(m ± sd) | DBP<br>(m ± sd) |
| --- | --- | --- | --- | --- | --- | --- |
| Females | 25 | 37.16 ± 12.6 [18.7-63.5] | 15.12 ± 1.9 | 69.36 ± 8.3 | 117.52 ± 16.6 | 72.96 ± 10.9 |
| Males | 22 | 38.69 ± 15.1 [19.8-69.0] | 15.41 ± 2.1 | 70.64 ± 11.6 | 118.86 ± 10.4 | 75.86 ± 5.3 |
| Total | 47 | 37.88 ± 13.7 [18.7-69.0] | 15.26 ± 2.0 | 69.96 ± 9.9 | 118.15 ± 13.9 | 74.32 ± 8.8 |

n, number of subjects; Age, decimal age; m, mean value, sd, standard deviation RR, respiratory rate; HR, heart rate; SBP, systolic blood pressure; DBP, diastolic blood pressure

The Ethics Committee of the Institute of Neurology and Neurosurgery approved this research and the participants gave their informed written consent to participate in the study. The clinical EEGs included a 10 minutes-long recording in a functional state of relaxed resting-state wakefulness with eyes-closed, and other physiological maneuvers. The EEG was recorded from 19 standard locations over the scalp according to the 10–20 system: Fp1, Fp2, F3, F4, F7, F8, T3, T4, C3, C4, P3, P4, T5, T6, O1, O2, Fz, Cz, and Pz. Silver scalp electrodes were fixed and connected to the input box of a digital EEG system (Medicid-05, Neuronic, S.A.). Monopolar leads were employed, using linked ears as a reference. EEG technical parameters were: gain 20,000, pass-band filters 0.5–70 Hz, ‘‘notch” filter at 60 Hz, noise level of 2 μV (root mean square), sampling frequency 200 Hz, A/D converter of 15 bits, and electrode–skin impedance never higher than 5 KΩ. Two of us (EAP and MEB) visually inspected the records to select free of artifacts EEG segments with a total duration of no less than 60 seconds, which were later exported to an ASCII file, for further quantitative analysis.

### 2.2 Pre-processing

A digital bandpass “FIR” filter created with Matlab using the function “desingnfilt.m” with cutoff frequencies fixed at 1 and 70 Hz was applied to the EEG 60-seconds segments digitally stored, using the “filtfilt.m” function. Finally, to avoid outliers, the “filloutliers.m” function was applied using for filling the outliers a piecewise cubic spline interpolation method.

### 2.3 PSD estimation using the traditional modified periodogram (FFT)

For the estimation of the PSD where used EEG windows of 1024 samples. For the 60-seconds EEG segments were detected 11 windows. For each window was calculated the power spectrum using the Matlab function “periodogram.m” with a Hamming window and 1024 FFT samples. The obtained spectra were then averaged to obtain the power spectrum of the studied EEG lead. The spectral resolution was 0.1953125 Hz.

### 2.4 Empirical mode decomposition (EMD)

EMD is a fully data-driven method for decomposing a time series into AM/FM components which reflect its natural oscillations. As EMD makes no prior assumptions on the data it becomes suitable for the analysis of nonlinear and non-stationary processes (Huang et al. 1998; Park et al. 2011). The EMD algorithm extracts the IMFs using an iterative technique called “sifting” (Huang et al. 1998). IMFs have two basic properties:

1. In the whole data set, the number of extrema is equal to the number of zero-crossings or differ at most by one.
2. At any point, the mean value of the envelope defined by the local maxima and the envelope defined by the local minima is zero.

Once an IMF is obtained, it is subtracted from the original signal and the procedure is repeated on the remaining signal to obtain the next IMF. The process is stopped when the signal at the end of an iteration becomes a constant, a monotonic function, or a function containing only a single extremum.

The signal x(t) can then be represented as in the expression

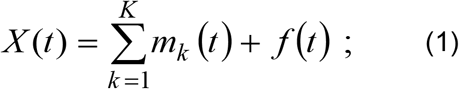

where m_k_ are the IMFs and f(t) is the final residual.

### 2.5 Multivariate empirical mode decomposition (MEMD)

Multivariate data, as is the case of the EEG signal, contain joint rotational modes whose coherent treatment is required for time-frequency estimation. The application of the algorithm of EMD to the EEG signals registered simultaneously from several areas of the scalp shows the presence of a different number of IMFs for the studied EEG leads (this is known as nonuniformity). Consequently, there isn’t guarantee that the same IMFs would contain equal scales across data channels. Besides, to enforce the same number of IMFs for every data channel may compromise time-frequency estimation, because such IMFs are typically not mono-component (Mandic et al. 2013).

In EMD, the local mean is calculated by taking an average of upper and lower envelopes, which in turn are obtained by interpolating between the local maxima and minima. However, generally, for multivariate signals, the local maxima and minima may not be defined directly. Besides, the notion of ‘oscillatory modes’ defining an IMF is rather confusing for multivariate signals. MEMD propose to generate multiple n-dimensional envelopes by taking signal projections along different directions in n-dimensional spaces; these are then averaged to obtain the local mean.

As the direction vectors in n-dimensional spaces can be equivalently represented as points on the corresponding unit (n - 1) spheres, the problem of finding a suitable set of direction vectors can be treated as that of finding a uniform sampling scheme on an n-sphere. A convenient choice for a set of direction vectors can be to employ uniform angular sampling of a unit sphere in an n-dimensional hyper-spherical coordinate system. A coordinate system in an n-dimensional Euclidean space can be defined to serve as a pointset (and the corresponding set of direction vectors) on an (n - 1) sphere To generate a uniform pointset on an n-sphere it is used the concept of discrepancy. A convenient method for generating multidimensional ‘low-discrepancy’ sequences involves the family of Halton and Hammersley sequences.

The algorithm of the MEMD (Rehman, Mandic 2010; Ur Rehman et al. 2010) include several steps that are briefly detailed in Table 2.

**Table 2.** Algorithm of the multivariate empirical mode decomposition.

| Main steps of the MEMD algorithm |
| --- |
| 1. Choose a suitable pointset for sampling on an $(n - 1)$ sphere. |
| 2. Calculate a projection of the input signal along the direction vector for the whole set of direction vectors giving the set of projections. |
| 3. Find the time instants corresponding to the maxima of the set of projected signals. |
| 4. Interpolate to obtain multivariate envelope curves. |
| 5. For a set of $K$ direction vectors, the mean $m(t)$ of the envelope curves is calculated as |
| $m(t) = \frac{1}{K} \sum_{k=1}^K e^{\theta_k}(t) ;$ |
| where $e^{\theta_k}$ are the envelope curves, and $K$ the direction vectors. |
| 6. Extract the “detail” $(t)$ using $(t) = (t) - (t)$. If the “detail” $(t)$ fulfills the stoppage criterion for a multivariate IMF, apply the above procedure to $(t) - (t)$, otherwise apply it to $(t)$. |

### 2.6 Adaptive-projection intrinsically transformed MEMD (APIT-MEMD)

Recently, it has been developed an extension to the MEMD to cater for power imbalances and inter-channel correlations in real-world multichannel data as the EEG (Hemakom et al. 2016). The main steps of this novel algorithm (Hemakom et al. 2016) are included in Table 3.

**Table 3.**
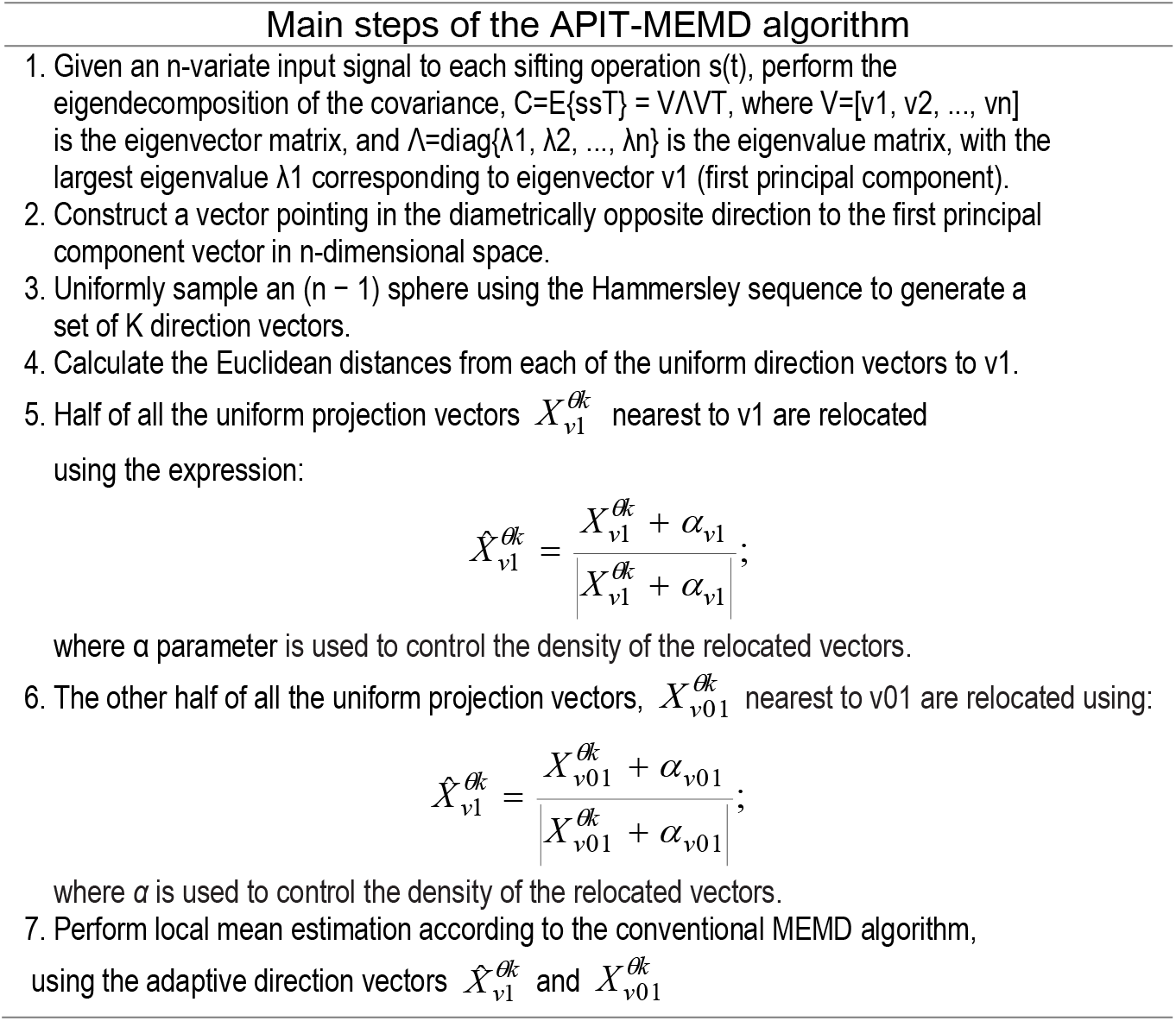
Algorithm of the adaptive-projection intrinsically transformed MEMD.

In this study we used the APIT-MEMD algorithm for the EEG decomposition, with an alpha parameter of 0.35 and the number of directions was of 128. As a rule of thumb, the minimum value of directions should be twice the number of data channels. In this study, for 19 channels, we used 128 directions that is at least six times the number of channels. The implementation of the MEMD and the APIT-MEMD algorithms used in this study was based on software resources publicly available at the web site http://www.commsp.ee.ic.ac.uk/~mandic/research/emd.htm. For calculations in the frequency domain of EEG signals were selected the first 6 extracted IMFs, that contain the spectral range from 1 to 70 Hz as shown by other authors (Chen et al. 2016; Chen et al. 2017; Schiecke et al. 2019; Amo et al. 2017; Carella et al. 2018; Zheng, Xu 2019; Zhuang et al. 2017; Tsai et al. 2016).

### 2.7 Hilbert marginal spectrum (HMS)

Each IMF extracted from the EEG was submitted to the Hilbert transformation. The corresponding instantaneous values of frequency, power and phase were then calculated. The application of the Hilbert transform to the original IMF allows the creation of an analytic signal referred here as Z(t). It can be written in this way

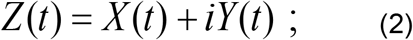

where X(t) is the input time series (the IMF in this case), and iY(t) is the Hilbert transform of X(t).

Several instantaneous indices of this analytic signal can now be obtained using the following expressions:

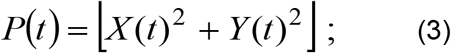

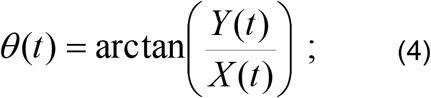

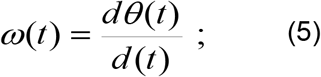

where P(t) is the instantaneous energy power, θ(t) is the instantaneous phase, and ω(t) is the instantaneous frequency (Estevez-Baez et al. 2019; Estevez-Baez et al. 2018). The instantaneous values of frequency and power for the EEG segments of 60 seconds were used for the calculation of the HMS. The spectral resolution used for the HMS was the same as the one for the traditional modified periodogram (0.1953125 Hz). In Figure 1 is presented a schematic diagram showing the main steps carried out for the processing of the data in this study.

**Figure 1.**
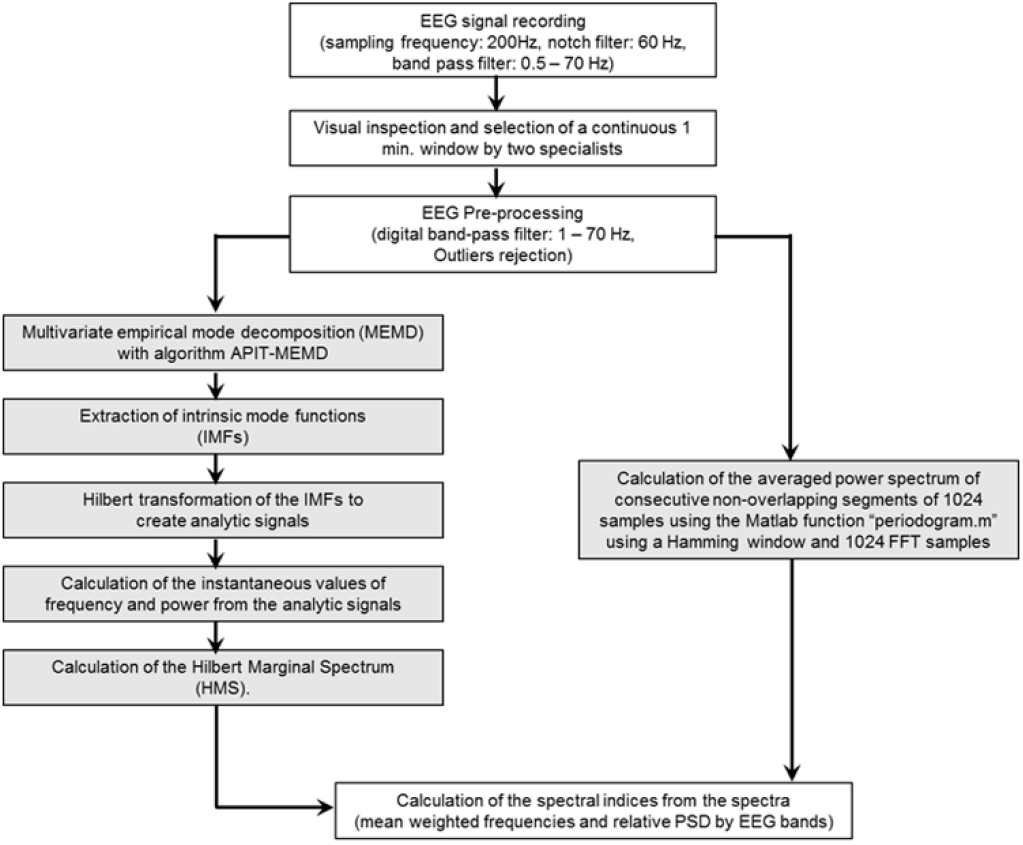
Block diagram of the main steps of the processing of the EEG used in this study.

### 2.8 Spectral indices

The absolute power values for each band were calculated following the expression:

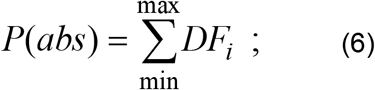

where ‘min’ represents the lower limit value of the spectral range of the band, ‘max’ the upper limit, and DFi are the discrete spectral frequencies of the spectra. The spectral relative power expressed in normalized units (%) was calculated as:

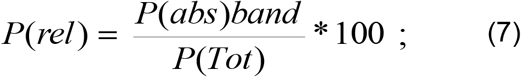

where P(abs)band is the power calculated for the band, and P(Tot) is the sum of the power of the six bands of the spectrum.

The mean weighted frequency (Xie, Wang 2006) of the spectral bands was calculated using the expression:

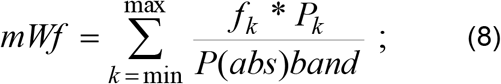

where f_k_ discrete spectral frequencies, P_k_ corresponding discrete spectral powers, P(abs)band is the sum of the absolute power of the band, “min” is the lower limit value of the spectral range of the band, and “max” the upper limit.

### 2.9 Statistical analysis

All results were calculated in this work using programs and functions developed with Matlab R2019b (9.7.0.1190202). It was explored for all the calculated indices the normality of their distribution using the Shapiro-Wilks, Lilliefors, and the Kolmogorov-Smirnov tests, and transformed when necessary to achieve a normal distribution before the statistical comparisons. The grand average method was applied for calculation of indices of mean differences of spectral frequencies, and to obtain average spectra for the whole group in different EEG leads. An ANOVA test of repeated measures was carried out for the comparison of the values obtained using both methods of spectral analysis using the statistical commercial package Statistica 10 (StatSoft, Inc. (2011).

## 3. Results

The empirical mode decomposition of the EEG using the multivariate APIT-MEMD algorithm showed in all cases for all the EEG leads the same number of extracted IMFs, with the faster oscillations for the IMF-1 and progressively slower in the consecutive IMFs. The decomposition of 2000 samples (10 seconds) of the first six IMFs of three EEG leads of one of the volunteers is shown, as an example in Figure 2. The instantaneous values of frequency and power obtained after the Hilbert transformation of the corresponding IMFs of an EEG segment of 2000 samples can be observed in Figure 3.

**Figure 2.**
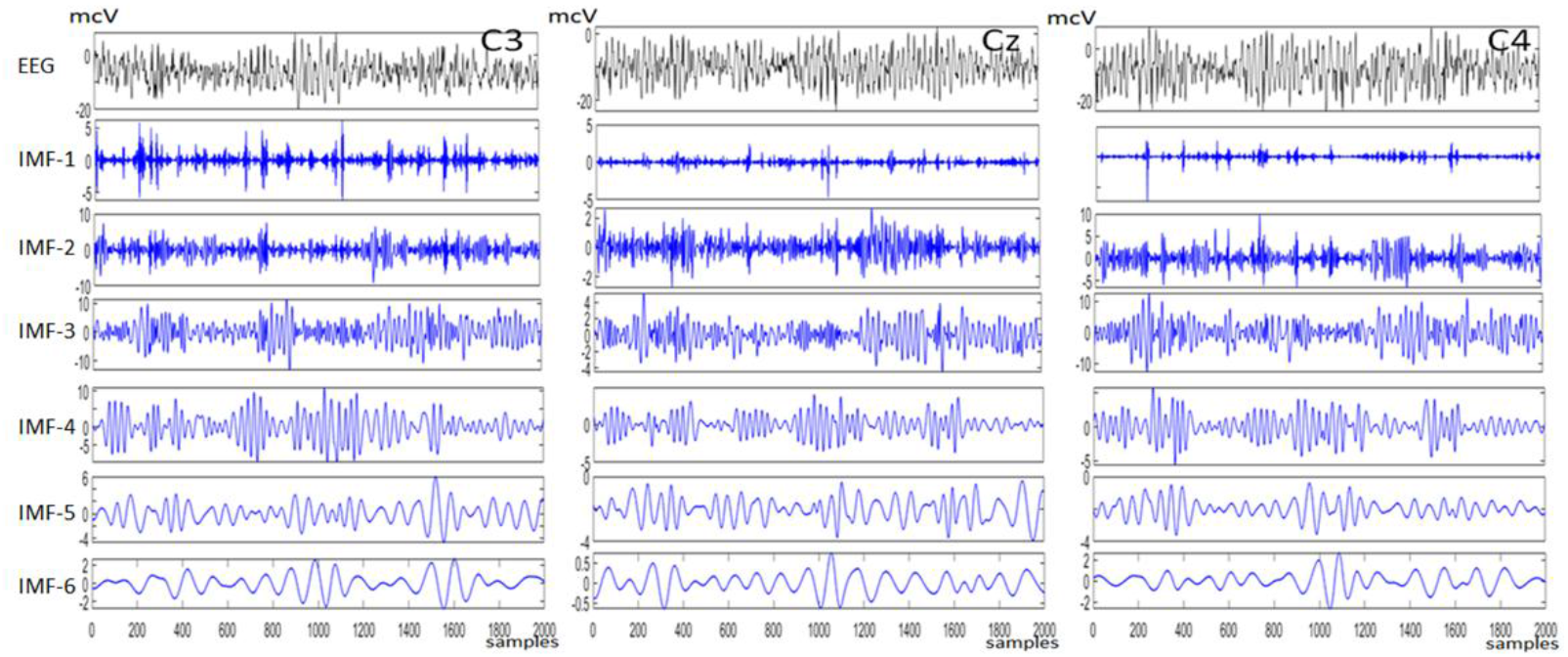
An example of decomposition using APIT-MEMD of 10 seconds-long EEG segments recorded at the C3, Cz, and C4 leads (International 10-20 System), in one of the healthy volunteers. The raw, original EEG, is shown at the top and below are depicted the first six consecutively extracted intrinsic mode functions (IMF).

**Figure 3.**
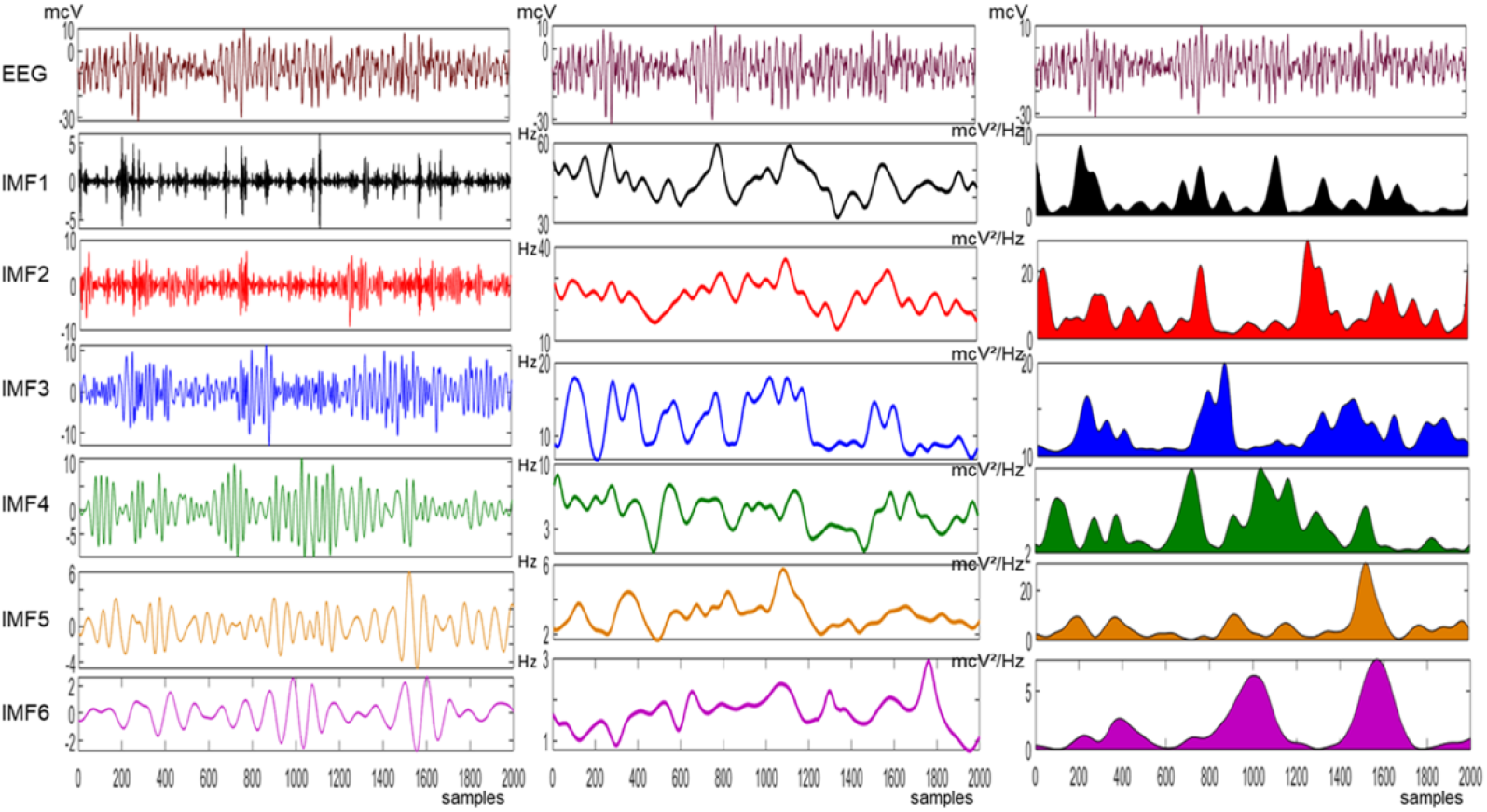
Collage showing the decomposition using the APIT-MEMD of a 10 seconds-long EEG segment (2000 samples) from the C3 EEG lead (left panel), and the corresponding instantaneous values of frequency (center panel) and power (right panel) for each extracted intrinsic mode functions (IMF-1 - IMF-6) obtained after application of the Hilbert transform to the extracted IMFs.

The values calculated with both methods (FFT and HMS) for the mean weighted frequency of the six EEG bands showed a normal distribution for all the EEG leads. However, the values calculated for the power spectral density (PSD) expressed in normalized units (%) with both methods didn’t show a normal distribution. In consequence, the original values were normalized using the “*logit*” transformation (log (x/1-x)) before the ANOVA test.

The values of the mean weighted frequency for the HMS were significantly lower than those observed with the traditional FFT spectrum for all the EEG leads for the delta band, and practically for all EEG leads for the theta band, and the beta-1 band. For the alpha, beta-2 and gamma bands the values were significantly higher for the HMS (See Table 4). A more detailed analysis of the differences found between mWf values is shown in Table 6. The grand average of the differences between mean values for all the EEG leads only resulted in higher or lower values from 0.16 to 1 Hz for the delta, theta, alpha, beta1 and beta2 bands, and only for the gamma band the values of the mean differences calculated indicated values of +3.4 Hz indicating higher values for the HMS method.

**Table 4.** Mean weighted frequency values (Hz) calculated for the EEG bands obtained from forty-seven healthy volunteers using two different methods for estimation of the PSD.

| Lead | Method | Delta | Theta | Alfa | Beta-1 | Beta-2 | Gamma |
| --- | --- | --- | --- | --- | --- | --- | --- |
| Fp1 | FFT | 2.77 ± 0.1 | 6.08 ± 0.3 | 9.80 ± 0.5 | 14.98 ± 0.2 | 22.48 ± 0.8 | 42.10 ± 3.0 |
|  | HMS | 2.64 ± 0.1 ↓ | 5.90 ± 0.3 ↓ | 9.98 ± 0.4 ↑ | 14.33 ± 0.2 ↓ | 23.46 ± 1.2 ↑ | 45.32 ± 3.5 ↑ |
| Fp2 | FFT | 2.75 ± 0.2 | 6.05 ± 0.3 | 9.81 ± 0.5 | 14.97 ± 0.2 | 22.40 ± 0.9 | 42.22 ± 3.3 |
|  | HMS | 2.63 ± 0.1 ↓ | 5.90 ± 0.3 ↓ | 9.99 ± 0.4 ↑ | 14.31 ± 0.2 ↓ | 23.33 ± 1.2 ↑ | 45.25 ± 3.8 ↑ |
| F3 | FFT | 2.85 ± 0.1 | 6.13 ± 0.3 | 9.80 ± 0.5 | 14.99 ± 0.2 | 22.29 ± 0.8 | 41.72 ± 2.9 |
|  | HMS | 2.68 ± 0.1 ↓ | 5.94 ± 0.3 ↓ | 9.98 ± 0.4 ↑ | 14.33 ± 0.2 ↓ | 23.33 ± 1.2 ↑ | 45.10 ± 3.2 ↑ |
| F4 | FFT | 2.86 ± 0.1 | 6.12 ± 0.3 | 9.82 ± 0.5 | 14.95 ± 0.2 | 22.28 ± 0.9 | 41.57 ± 2.5 |
|  | HMS | 2.68 ± 0.1 ↓ | 5.95 ± 0.3 ↓ | 10.01 ± 0.4 ↑ | 14.34 ± 0.2 ↓ | 23.19 ± 1.3 ↑ | 44.89 ± 3.0 ↑ |
| C3 | FFT | 2.89 ± 0.1 | 6.19 ± 0.3 | 9.86 ± 0.5 | 14.97 ± 0.2 | 22.04 ± 0.9 | 41.38 ± 3.0 |
|  | HMS | 2.69 ± 0.1 ↓ | 6.03 ± 0.3 ↓ | 10.00 ± 0.4 ↑ | 14.33 ± 0.2 ↓ | 23.02 ± 1.3 ↑ | 44.83 ± 3.2 ↑ |
| C4 | FFT | 2.89 ± 0.1 | 6.19 ± 0.3 | 9.86 ± 0.5 | 14.97 ± 0.2 | 21.98 ± 0.9 | 41.51 ± 3.2 |
|  | HMS | 2.70 ± 0.1 ↓ | 6.02 ± 0.2 ↓ | 10.00 ± 0.4 ↑ | 14.32 ± 0.2 ↓ | 23.03 ± 1.3 ↑ | 45.10 ± 3.4 ↑ |
| P3 | FFT | 2.87 ± 0.1 | 6.25 ± 0.4 | 9.91 ± 0.5 | 14.95 ± 0.2 | 21.75 ± 0.9 | 41.40 ± 3.3 |
|  | HMS | 2.70 ± 0.1 ↓ | 6.09 ± 0.3 ↓ | 10.00 ± 0.4 | 14.32 ± 0.2 ↓ | 22.73 ± 1.3 ↑ | 45.13 ± 3.5 ↑ |
| P4 | FFT | 2.88 ± 0.1 | 6.21 ± 0.3 | 9.89 ± 0.5 | 14.95 ± 0.2 | 21.73 ± 0.9 | 41.16 ± 3.1 |
|  | HMS | 2.71 ± 0.1 ↓ | 6.06 ± 0.3 ↓ | 9.98 ± 0.4 | 14.33 ± 0.2 ↓ | 22.71 ± 1.3 ↑ | 45.08 ± 3.4 ↑ |
| O1 | FFT | 2.88 ± 0.1 | 6.15 ± 0.3 | 9.90 ± 0.5 | 14.90 ± 0.2 | 21.72 ± 0.9 | 41.77 ± 3.1 |
|  | HMS | 2.70 ± 0.1 ↓ | 6.08 ± 0.3 | 9.95 ± 0.5 | 14.27 ± 0.2 ↓ | 22.75 ± 1.3 ↑ | 45.28 ± 3.5 ↑ |
| O2 | FFT | 2.89 ± 0.1 | 6.18 ± 0.3 | 9.88 ± 0.6 | 14.90 ± 0.2 | 21.71 ± 0.9 | 41.59 ± 2.9 |
|  | HMS | 2.69 ± 0.1 ↓ | 6.11 ± 0.3 | 9.93 ± 0.5 | 14.32 ± 0.2 ↓ | 22.73 ± 1.4 ↑ | 45.08 ± 3.1 ↑ |
| F7 | FFT | 2.83 ± 0.1 | 6.08 ± 0.3 | 9.78 ± 0.5 | 14.97 ± 0.2 | 22.31 ± 0.8 | 41.94 ± 2.1 |
|  | HMS | 2.67 ± 0.1 ↓ | 5.92 ± 0.3 ↓ | 9.96 ± 0.4 ↑ | 14.33 ± 0.2 ↓ | 23.32 ± 1.3 ↑ | 45.02 ± 2.8 ↑ |
| F8 | FFT | 2.85 ± 0.1 | 6.07 ± 0.3 | 9.79 ± 0.5 | 14.95 ± 0.2 | 22.25 ± 0.8 | 42.37 ± 2.6 |
|  | HMS | 2.68 ± 0.1 ↓ | 5.92 ± 0.3 ↓ | 9.98 ± 0.4 ↑ | 14.32 ± 0.2 ↓ | 23.33 ± 1.3 ↑ | 45.42 ± 3.3 ↑ |
| T3 | FFT | 2.91 ± 0.1 | 6.15 ± 0.3 | 9.84 ± 0.5 | 14.96 ± 0.2 | 22.21 ± 0.8 | 42.15 ± 2.3 |
|  | HMS | 2.68 ± 0.1 ↓ | 5.97 ± 0.3 ↓ | 10.02 ± 0.4 ↑ | 14.33 ± 0.2 ↓ | 23.31 ± 1.3 ↑ | 45.10 ± 2.9 ↑ |
| T4 | FFT | 2.90 ± 0.1 | 6.14 ± 0.3 | 9.84 ± 0.5 | 14.95 ± 0.2 | 22.20 ± 0.9 | 42.38 ± 2.7 |
|  | HMS | 2.71 ± 0.1 ↓ | 5.96 ± 0.3 ↓ | 10.00 ± 0.4 ↑ | 14.35 ± 0.2 ↓ | 23.23 ± 1.4 ↑ | 45.30 ± 3.2 ↑ |
| T5 | FFT | 2.88 ± 0.1 | 6.21 ± 0.4 | 9.79 ± 0.5 | 14.95 ± 0.2 | 21.92 ± 0.9 | 42.09 ± 2.7 |
|  | HMS | 2.68 ± 0.1 ↓ | 6.04 ± 0.3 ↓ | 9.92 ± 0.5 ↑ | 14.28 ± 0.2 ↓ | 23.05 ± 1.3 ↑ | 45.26 ± 3.4 ↑ |
| T6 | FFT | 2.89 ± 0.1 | 6.22 ± 0.3 | 9.79 ± 0.5 | 14.94 ± 0.2 | 21.82 ± 0.9 | 41.97 ± 2.9 |
|  | HMS | 2.70 ± 0.1 ↓ | 6.02 ± 0.3 ↓ | 9.92 ± 0.4 ↑ | 14.33 ± 0.2 ↓ | 22.94 ± 1.4 ↑ | 45.32 ± 3.2 ↑ |
| Fz | FFT | 2.86 ± 0.1 | 6.12 ± 0.3 | 9.80 ± 0.5 | 14.97 ± 0.2 | 22.20 ± 0.9 | 41.43 ± 3.1 |
|  | HMS | 2.70 ± 0.1 ↓ | 5.95 ± 0.3 ↓ | 9.98 ± 0.4 ↑ | 14.34 ± 0.2 ↓ | 23.19 ± 1.2 ↑ | 45.08 ± 3.4 ↑ |
| Cz | FFT | 2.87 ± 0.1 | 6.14 ± 0.3 | 9.82 ± 0.5 | 14.96 ± 0.2 | 22.19 ± 0.9 | 40.58 ± 2.1 |
|  | HMS | 2.71 ± 0.1 ↓ | 5.99 ± 0.3 ↓ | 9.97 ± 0.4 ↑ | 14.32 ± 0.2 ↓ | 23.11 ± 1.2 ↑ | 44.43 ± 2.8 ↑ |
| Pz | FFT | 2.87 ± 0.1 | 6.20 ± 0.3 | 9.88 ± 0.5 | 14.94 ± 0.3 | 21.83 ± 0.9 | 40.87 ± 2.2 |
|  | HMS | 2.70 ± 0.1 ↓ | 6.07 ± 0.3 ↓ | 9.97 ± 0.4 ↑ | 14.31 ± 0.2 ↓ | 22.73 ± 1.2 ↑ | 44.73 ± 2.8 ↑ |

FFT, spectral method based on the Fast Fourier Transform; HMS, spectral method based on the Hilbert Marginal Spectrum; ↑\_ Significant higher, or ↓\_ lower values (p < 0.05) for the ANOVA "F" statistics.

The values of the PSD were significantly higher for most the EEG leads in the delta, theta, beta-1 and gamma bands calculated with the HMS method compared with those of the traditional FFT spectrum. Values in the alpha band showed significant reduced values, while for the beta-2 band were not found differences between methods. (See Table 5).

**Table 5.**
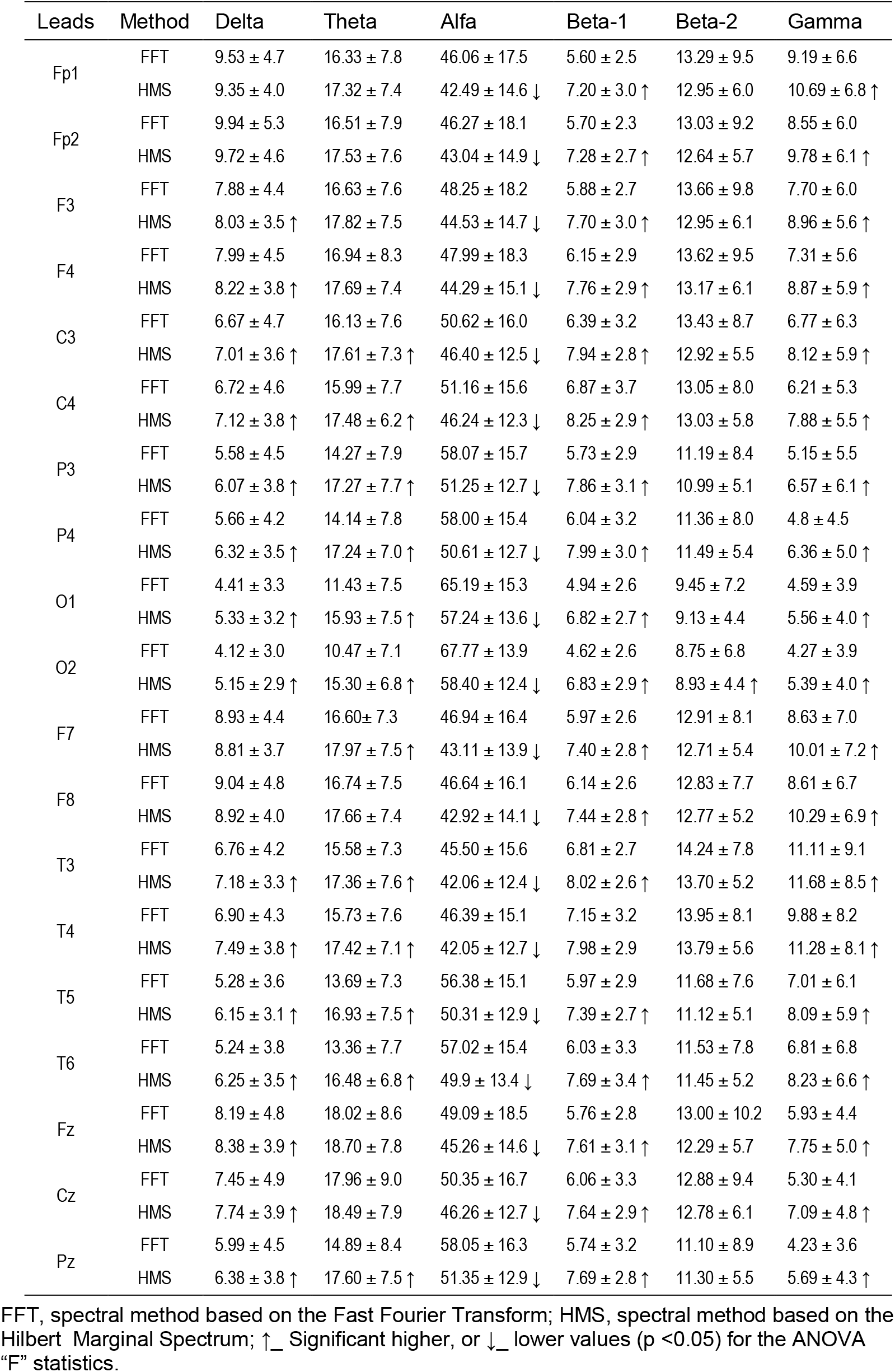
Power spectral density expressed in normalized units (%) calculated for the EEG bands obtained from forty-seven healthy volunteers using the FFT and the HMS methods.

The analysis using grand averages of the differences of mean values calculated in all the EEG leads (Table 6) evidenced that the reduction in relative PSD observed for the alpha band was of approximately 5.6%, and was the result of a corresponding increment in other bands that in decreasing order of magnitude were 2.13% for the theta band, 1.67% for beta-1, 1.38% for gamma band, and 0.45% for the delta band, not showing differences for the beta-2 band.

**Table 6.** Grand average of significant mean values changes calculated from the differences between methods (HMS-FFT) considering all EEG leads for both spectral indices and to each the band.

| Spectral Indices | Delta (m ± sd) | Theta (m ± sd) | Alfa (m ± sd) | Beta-1 (m ± sd) | Beta-2 (m ± sd) | Gamma (m ± sd) |
| --- | --- | --- | --- | --- | --- | --- |
| mWf | -0.177 ± 0.03 | -0.165 ± 0.02 | +0.161 ± 0.03 | -0.631 ± 0.02 | +1.011 ± 0.07 | +3.395 ± 0.32 |
| PSD nU | +0.454 ± 0.31 | +2.139 ± 1.32 | -5.641 ± 1.86 | +1.674 ± 0.28 | -- | +1.381 ± 0.30 |

m, mean value; sd, standard deviation; mWf, mean weighted frequency; *PSD nU*, power spectral density expressed in normalized units (%) of the total energy; + or -, indicates the sense of the calculated grand average of the differences calculated as "values HMS – values FFT"; -- values not calculated because there were not found significant differences for values in the band for any EEG lead.

In Figures 4 and 5 are presented the power spectra for the 19 EEG leads obtained with the traditional FFT spectrum method, and the HMS in one of the healthy volunteers. The grand averaged spectra of EEG leads for the whole group of healthy volunteers allowed to visually confirm the statistical results presented in Table 5.

**Figure 4.**
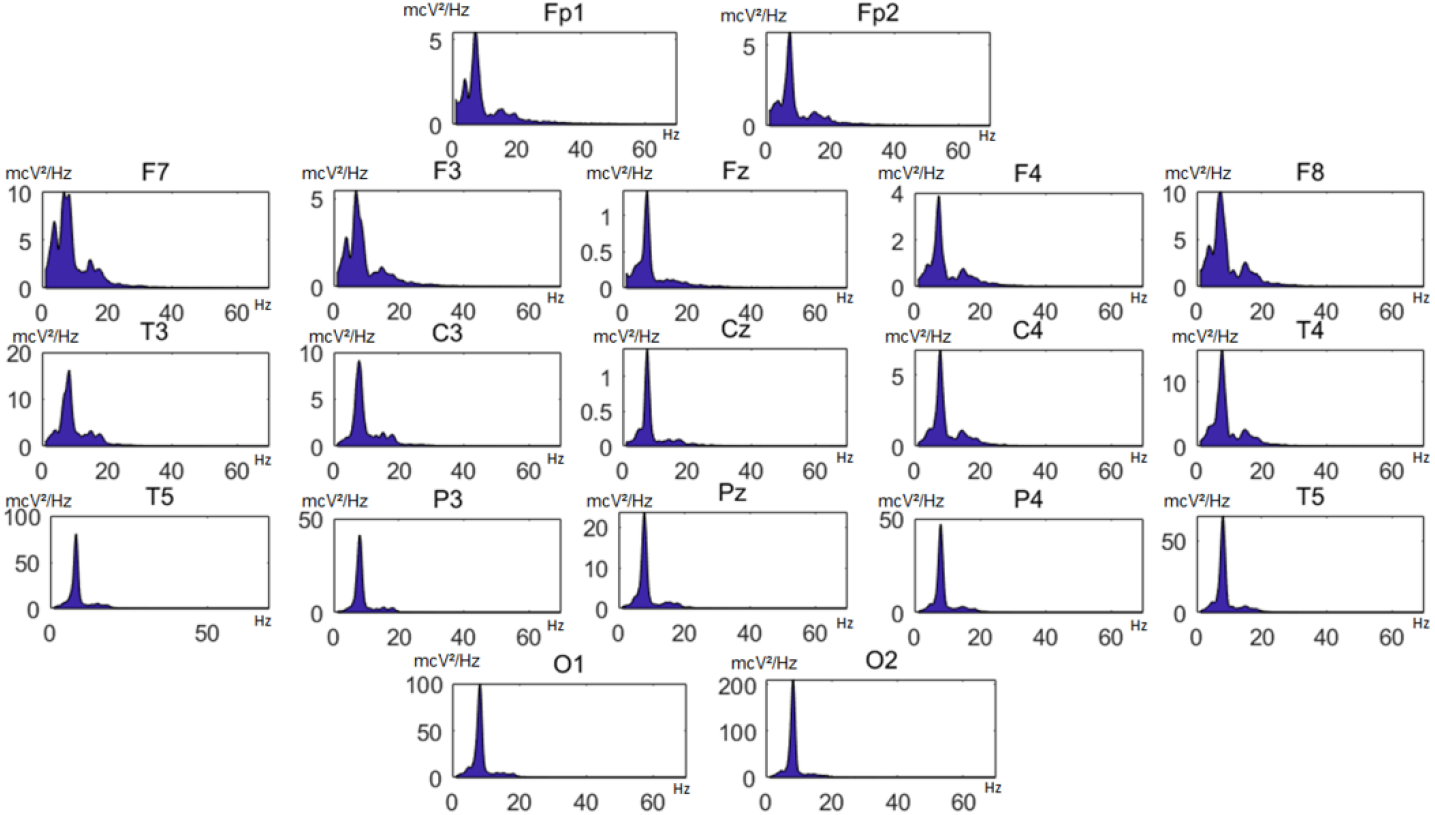
Power spectra calculated with the traditional modified periodogram FFT method in a representative subject of the group of studied healthy volunteers; for visual purposes in the graphic, a sliding smoothing window with a length 5 was applied to the original values of the power spectra.

**Figure 5.**
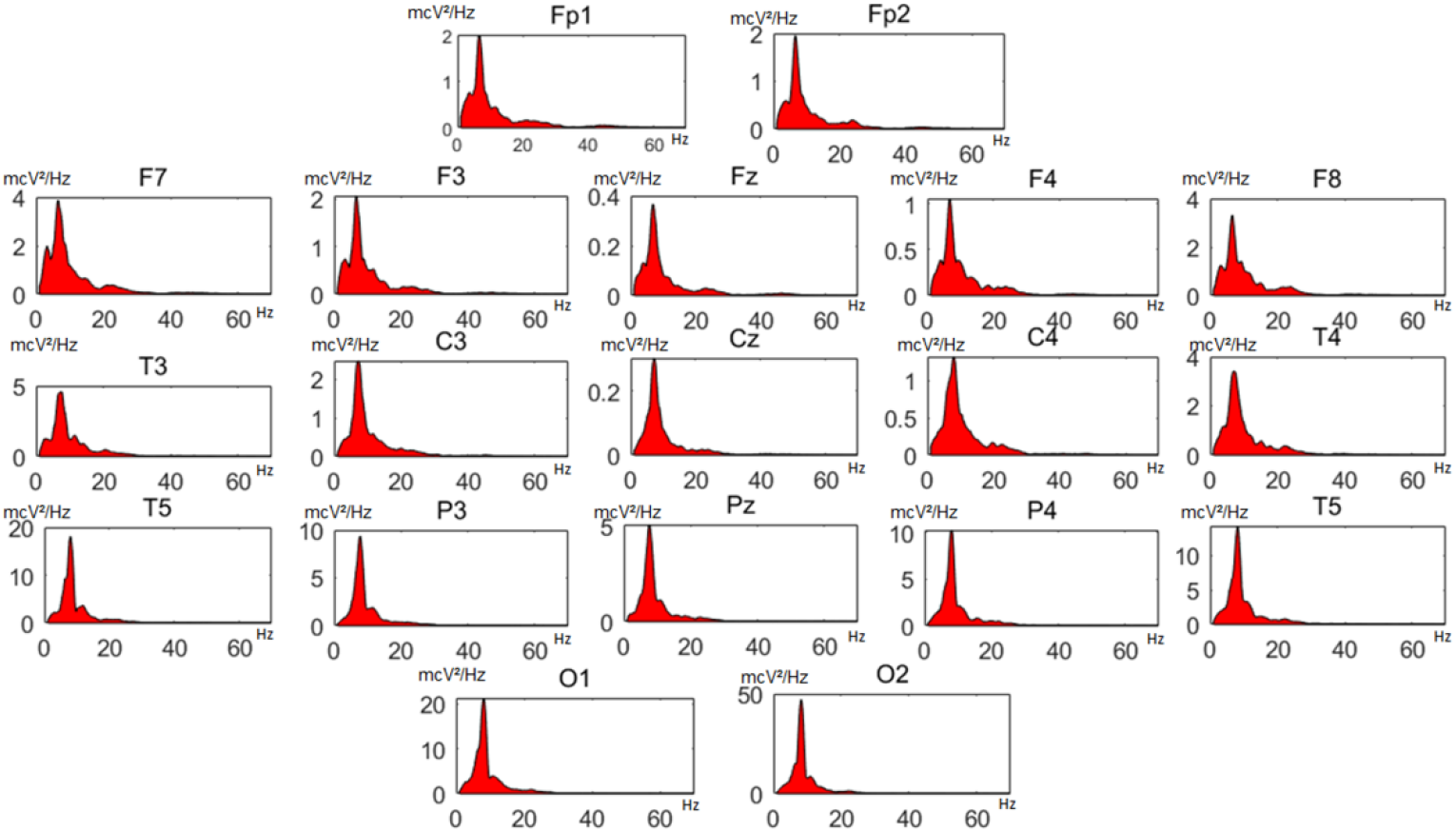
Power spectra calculated with the Hilbert marginal method in the same healthy volunteer as in Figure 4. exclusively for visual purposes in the graphic, a sliding smoothing window with a length 5 was applied to the original values of the power HMS spectra.

The grand averages of the spectra for the EEG leads F3, C3, T3, and O1, calculated with both methods, as an example of the general findings in the studied group, are depicted in Figure 6.

**Figure 6.**
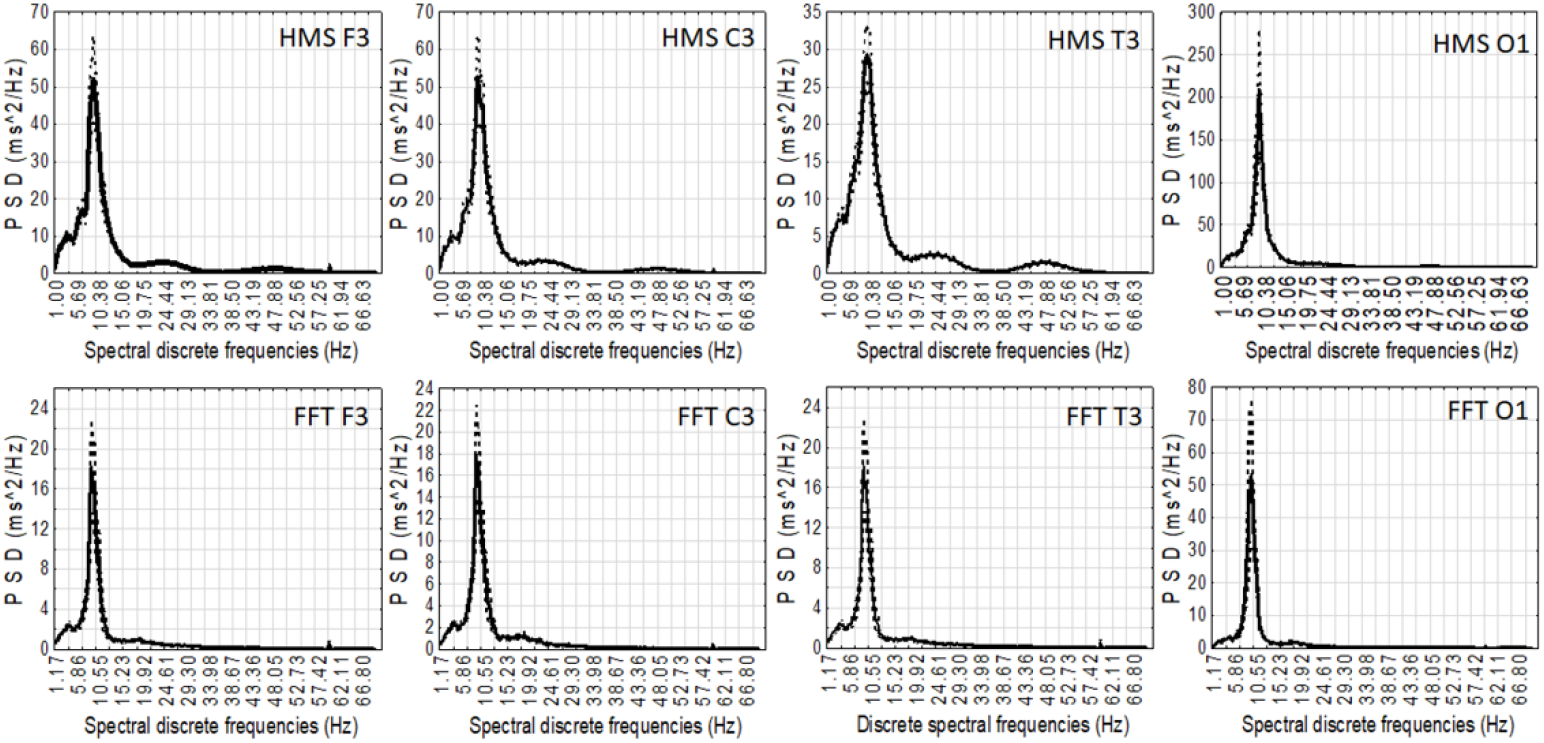
Grand average of the spectra obtained for some EEG leads using the traditional FFT method and the HMS in 47 healthy volunteers. The bold lines depict the mean grand average, and the intermittent lines depict the values of ± 1 standard error of the mean. P.S.D: power spectral density; no smoothing was used to calculate the grand averages nor to the HMS spectra included in this particular graphic.

## 4. Discussion

Spectral indices of the broadband analysis of the EEG in 6 frequency bands showed significant statistical differences between values calculated with both methods, but a detailed analysis of these differences showed that the two methods estimated the spectral parameters with an enough similarity for practical purposes. Nevertheless, it results useful to analyze the detected statistical differences because they reflect particularities associated with each method of analysis.

Slightly reduced values of the relative PSD expressed in normalized units (%) in the alpha range considered in this study (8-13 hz), observed for the marginal spectrum method, were the result of a better detection using this technique of higher values in other ranges of the spectra, and particularly the increase in the beta1 and gamma fast frequency bands, and in the theta band. In the last years the study of the beta band EEG activity has been the object of motor imagery classification study for brain computer interfaces and assessment of movement of the fingers (Chang et al. 2013; Kim et al. 2016; Park et al. 2013) using the EMD and the MEMD, and it has been shown that the spectra of the IMFs corresponding to the beta frequencies give a better quantitative information than the FFT method applied directly to the EEG original signal. This fact has been also confirmed for the detection of gamma band activity during motor tasks (Amo et al. 2017), to assess the gamma activity in the coordination of spatially directed limb and eye movements (Park et al. 2014), and to demonstrate the increase of the gamma band power triggered by the anesthetic Ketamine (Tsai et al. 2016). These results agree with the observed increments in beta-1 and gamma bands of the HMS observed in our present study.

The slight but significant differences observed for values of the mean weighted frequency in the different bands may be associated with the ability of the Hilbert-Huang method that enhances the presence of fast low amplitude oscillations, and offers better time-resolution due to its instantaneous frequency property (Munoz-Gutierrez et al. 2018), and performs better compared with Fourier analysis (Noshadi et al. 2014).

Even though the HMS has been used by other authors for particular applications on EEG spectral analysis (Chen et al. 2010; Chen et al. 2016; Li 2006; Li et al. 2008; Xiangjun et al. 2017; Zhu et al. 2015; Park et al. 2011) the possible differences with the traditional FFT spectrum have not been analyzed in details particularly in healthy humans, as we have described in this study, and could be considered the first report about this particular issue. The fact that we have shown no substantial differences for the typical indices calculated for broadband EEG spectral analysis using both methods in this study, can be considered a good evidence that the use of the classic FFT method, applied extensively and for many years, can be used in the conditions in which was carried out this investigation, it is, in healthy subjects, during a functional state of relaxed wakefulness during resting eyes-closed condition, and for segments of consecutive EEG of 60 seconds, but for other physiological conditions, during cognitive studies, or to study patients with different pathologies that can affect the processes responsible of the generation of the bioelectric activity of the brain, the presence of nonlinearity and particularly of non-stationarity may produce a misleading result while using the FFT methods (Alegre-Cortes et al. 2016; Huang et al. 1998; Munoz-Gutierrez et al. 2018; Soler et al. 2020; Tsai et al. 2016). Then, an alternative could be the use of the marginal spectrum after extracting the IMFs using an appropriate method of empirical mode decomposition, as has been described in this investigation.

Recently, we have shown that the use of the HMS applied to the heart rate variability analysis can outperform the traditional FFT method for the assessment of patients with cardiac autonomic neuropathy (Estevez-Baez et al. 2018) and in patients in coma (Estevez-Baez et al. 2019). The results shown in this study strongly support its introduction as an alternative to the traditional FFT spectrum for the spectral analysis of the EEG considering that the Hilbert-Huang method has been developed for the analysis of nonlinear and non-stationary signals.

However, the APIT-multivariate empirical mode decomposition algorithm used in this study (Hemakom et al. 2016) using 128 directions to obtain the intrinsic mode functions is a highly demanding computing task. For the study of a 19-leads EEG it is time consuming and for more EEG leads it can only be possible using powerful and fast processors. But undoubtedly, its performance is superior to other multivariate methods as the MEMD, and of course to the univariate EMD method, that must be avoided for the study of EEG signals recorded from different locations of the scalp simultaneously.

## 5. Conclusions

The traditional FFT method for spectral analysis of the EEG and the Hilbert marginal spectra calculated with a multivariate empirical mode decomposition for 60-seconds’ duration continuous EEG segments of healthy humans during a resting wakeful closed-eyes physiological condition show substantial similarity for the most frequently explored indices of frequency and relative PSD used in typical broadband analysis. Particular differences observed between the methods may be related with known abilities of the enhancement of fast low amplitude oscillations, and a better time-resolution of the Hilbert marginal spectra due to its instantaneous frequency property, that allows a better performance than the classical FFT approach. In those experimental conditions where nonlinear and/or non-stationarity may be present in the EEG, the alternative use of the marginal spectrum after the Hilbert-Huang transform is highly recommended to avoid possible misleading results. The marginal spectrum seems particularly useful for those studies of low amplitude fast frequency oscillations in the EEG bands beta and gamma.

## Competing Interests

The authors declare that they have no competing interests in relation to this article.

## Notes

### Competing Interest Statement

The authors have declared no competing interest.

